# Geographic shifts in *Aedes aegypti* habitat suitability in Ecuador using larval surveillance data and ecological niche modeling: Implications of climate change for public health vector control

**DOI:** 10.1101/404293

**Authors:** Catherine A. Lippi, Anna M. Stewart-Ibarra, M.E. Franklin Bajaña Loor, Jose E. Dueñas Zambrano, Nelson A. Espinoza Lopez, Jason K. Blackburn, Sadie J. Ryan

## Abstract

Arboviral disease transmission by *Aedes* mosquitoes poses a major challenge to public health systems in Ecuador, where constraints on health services and resource allocation call for spatially informed management decisions. Employing a unique dataset of larval occurrence records provided by the Ecuadorian Ministry of Health, we used ecological niche models (ENMs) to estimate the current geographic distribution of *Aedes aegypti* in Ecuador, using mosquito presence as a proxy for risk of disease transmission. ENMs built with the Genetic Algorithm for Rule-Set Production (GARP) algorithm and a suite of environmental variables were assessed for agreement and accuracy. The top model of larval mosquito presence was projected to the year 2050 under various combinations of greenhouse gas emissions scenarios and models of climate change. Under current climatic conditions, larval mosquitoes were not predicted in areas of high elevation in Ecuador, such as the Andes mountain range, as well as the eastern portion of the Amazon basin. However, all models projected to scenarios of future climate change demonstrated potential shifts in mosquito distribution, wherein range contractions were seen throughout most of eastern Ecuador, and areas of transitional elevation became suitable for mosquito presence. Encroachment of *Ae. aegypti* into mountainous terrain was estimated to affect up to 4,215 km^2^ under the most extreme scenario of climate change, an area which would put over 12,000 people currently living in transitional areas at risk. This distributional shift into communities at higher elevations indicates an area of concern for public health agencies, as targeted interventions may be needed to protect vulnerable populations with limited prior exposure to mosquito-borne diseases. Ultimately, the results of this study serve as a tool for informing public health policy and mosquito abatement strategies in Ecuador.

**Author summary:** The yellow fever mosquito (*Aedes aegypti*) is a medically important vector of arboviral diseases in Ecuador, such as dengue fever and chikungunya. Managing *Ae. aegypti* is a challenge to public health agencies in Latin America, where the use of limited resources must be planned in an efficient, targeted manner. The spatial distribution of *Ae. aegypti* can be used as a proxy for risk of disease exposure, guiding policy formation and decision-making. We used ecological niche models in this study to predict the range of *Ae. aegypti* in Ecuador, based on agency larval mosquito surveillance records and layers of environmental predictors (e.g. climate, altitude, and human population). The best models of current range were then projected to the year 2050 under a variety of greenhouse gas emissions scenarios and models of climate change. All modeled future scenarios predicted shifts in the range of *Ae. aegypti*, allowing us to assess human populations that may be at risk of becoming exposed to *Aedes* vectored diseases. As climate changes, we predict that communities living in areas of transitional elevation along the Andes mountain range are vulnerable to the expansion of *Aedes aegypti*.

## Introduction

Mosquito-borne disease transmission poses an ongoing challenge to global public health. This is especially true in much of Latin America, where arboviral disease management is complicated by the proliferation of mosquito vectors in tropical conditions, frequently coupled with limited resources for medical care and comprehensive vector control services [1]. In Ecuador, the *Aedes aegypti* mosquito is of particular medical importance as it is a competent vector for several established and emerging viral diseases, including all four serotypes of dengue virus (DENV), chikungunya (CHKV), Zika virus (ZKV), and yellow fever virus (YFV) [2–5]. The *Ae. albopictus* mosquito, also a competent vector of arboviruses, was recently reported for the first time in the city of Guayaquil, Ecuador [6]. Many of the diseases transmitted by *Ae. spp.* have no treatment beyond palliative care, and with the exception of yellow fever and dengue fever, there are no clinically established vaccines [7–9]. As a result, mosquito surveillance and control remain the best tools available for preventing and managing outbreaks of arboviral disease.

In Ecuador, the Ministry of Health (Ministerio de Salud Pública (MSP)) oversees public health vector control services in the country, including mosquito surveillance, indoor residual spraying, larvicide application, and ultra-low volume (ULV) fogging. The MSP conducts larval index (LI) surveys at the household level, wherein containers of impounded water are sampled for mosquito larvae. Larval indices are among the most common indicators used by public health agencies to establish mosquito presence and quantify abundance, key considerations for understanding localized transmission potential and planning larval source reduction [10]. Although cost effective relative to the delivery of clinical services, mosquito abatement and surveillance activities are nevertheless limited by financial constraints, necessitating informed strategies for focusing resources and personnel [11,12]. This becomes a critical factor when developing surveillance and control programs on very large scales, such as an entire country, where misdirection of program activities can rapidly deplete program funding. Advancing the understanding of where vectors of interest can occur on the landscape would provide valuable guidance in communicating risk of exposure and avoiding the pitfalls associated with indiscriminately rolling out interventions.

Like many mosquito species, the presence of *Ae. spp.* on the landscape is closely tied to environmental conditions [13,14]. Adult survival and larval development are largely driven and restricted by temperature, while successful oviposition and larval emergence rely on the persistence of impounded water in the environment [15–20]. In contrast with other medically important mosquito species in the region, such as Anopheline vectors of malaria, *Ae. aegypti* typically does very well in heavily urbanized environments, largely due to their reproductive strategy of exploiting small volumes of water in containers around the home as larval habitat [21]. In landscapes with heterogeneous topography, elevation also serves as a limiting factor for mosquito distributions, as temperature and precipitation change with altitude [22]. Situated in northwestern South America, Ecuador exemplifies high landscape diversity, with hot, humid areas of low elevation along the Pacific coast in the west and interior Amazon basin in the east, and the cool, arid Andes mountain range in the central portion of the country (Fig 1). Historically, the western coastal and interior regions experience a much higher incidence in mosquito-borne diseases than mountainous areas, where sharp increases in elevation and decreases in temperature limit the geographic distribution and vectorial capacity of the mosquito vector.

**Fig. 1.**
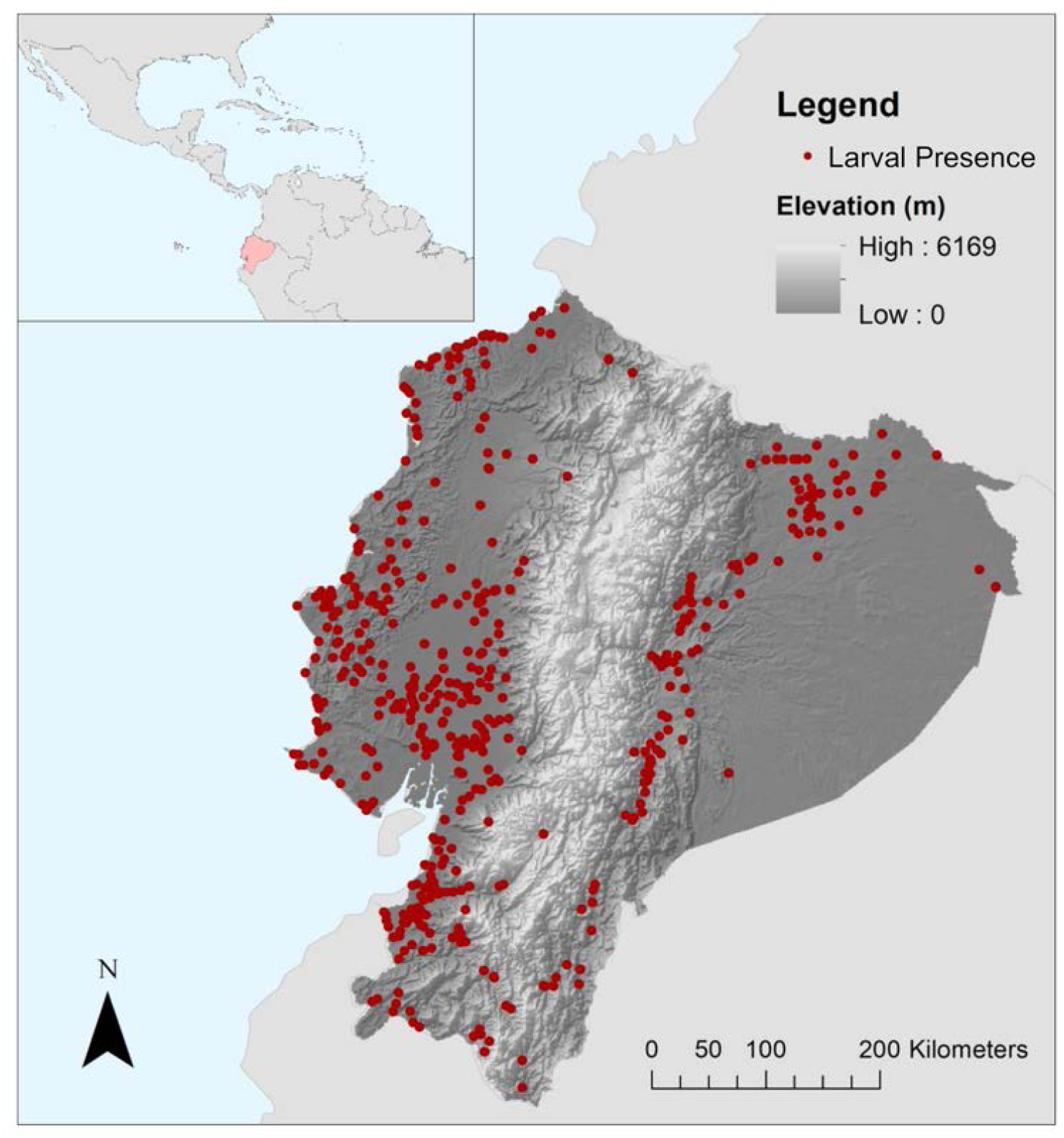
Ecuador, situated on the northwestern coast of South America (inset), has historically high prevalence of mosquito-borne diseases. The Ecuadorian Ministerio de Salud Pública (MSP) conducted household entomological surveys of *Aedes aegypti* throughout the country from 2000 – 2012. Spatially unique larval index (LI) occurrence records (n=478) collected in the survey were aggregated to cities and towns and used to model the ecological distribution of *Ae. aegypti* in Ecuador.

The present-day distribution of *Ae. aegypti* is broadly defined by regional temperature and precipitation trends, but global climate change has the potential to significantly alter the future geographic range of mosquito vectors [3]. The Intergovernmental Panel on Climate Change has established four representative concentration pathways (RCP), or different scenarios for future greenhouse gas emissions, which are the basis for modeling future climates. Even under the most conservative of these scenarios (RCP 2.6), mean global temperatures are projected to increase [23]. As temperature trends increase globally, it has been estimated that observed patterns in the distribution of mosquito vectors will shift accordingly; previous studies have projected that *Aedes* mosquitoes will increase their global range as temperature and rainfall patterns become more suitable under various climate change scenarios [17,24–26]. Modeling and visualizing changes in mosquito distributions at the national level will provide a useful tool for managing disease and planning the delivery of health services, as public health resources can be better allocated in anticipation of disease emergence in naïve populations driven by mosquito range expansions.

Ecological niche models (ENMs) have been used to estimate current potential distributions in insect populations, including mosquitoes, as well as range expansions resulting from environmental and climate changes [27–30]. ENM methodologies have been applied to many systems, spanning regional to global scales, in an effort to estimate *Aedes aegypti* distribution and the associated risk of exposure to humans, often indicating that water availability and land cover factor heavily into overall mosquito habitat suitability [3,27,31,32]. In Ecuador and other areas of Latin America, elevation also becomes a limiting factor for *Ae. aegypti* presence, though it is suggested that climate change may allow for the encroachment of mosquitoes into higher elevations [30,33]. While many prior studies have utilized records of adult stages of mosquitoes for ENMs, this study leverages existing larval surveillance data collected in Ecuador, providing a predictive tool about the source of mosquitoes in the environment. This complements predictive models for adult stages, particularly in considering potential for intervention, as it can target larvicidal approaches, rather than reactive adulticidal spraying methods. The Genetic Algorithm for Rule-Set Production (GARP) is a machine-learning algorithm that builds species ENMs using presence-only occurrence records and continuous environmental variables [34]. The genetic algorithm (GA) employed by GARP to build rule-sets for distribution models is stochastic in nature, resulting in a set of models from a single dataset of species occurrence records and allowing for the assessment of agreement between resulting models. This methodology offers a robust option for modeling the potential distribution of species on a landscape from presence-only records, as absence of a species is difficult to discern through historical records and field sampling (e.g. entomological surveys) [34,35]. GARP also provides a platform for projecting future climate scenarios onto the landscape with the natively generated rule-sets for species distribution prediction, allowing for the estimation of future geographic distributions [36].

Assessing current and future vector distributions in an ENM framework is useful for defining the spatial distribution and possible changes in risk exposure, using mosquito presence as a proxy for transmission risk. Previous work in Ecuador’s southern coast has focused on describing interannual variation in dengue transmission for a single region [37,38]. Here, we advance climate services available to the public health sector in Ecuador by providing climate-informed tools to assist decision-making, examining potential geographic shifts in risk at broader spatial and temporal scales. In this study, we had three main objectives 1) use an ENM approach to estimate the current geographic range of *Aedes aegypti* in Ecuador using a unique set of larval survey data; 2) use projected climate data to model the future geographic range under a variety of climate change scenarios; and 3) compare current and future climate models to describe changes in *Ae. aegypti* range over time, where we hypothesized that larval *Ae. aegypti* distribution in Ecuador would expand into areas of higher elevation with projected increases in global temperature.

## Methods

### Data sources

From 2000 – 2012 the MSP sampled aquatic larval mosquitoes from impounded water in and around households, in cities and towns throughout mainland Ecuador. These data were collected year-round by vector control technicians from the National Service for the Control of Vector-Borne Diseases (SNEM) of the MSP as part of routine vector surveillance activities. Positive LI records for *Aedes aegypti* were de-identified and aggregated to the administrative level of parroquia (township or parish) by the MSP for each year of the study. These de-identified, spatially aggregated data were made available to this study by the MSP.

### Informed disaggregation

Parroquias represented in this data set range in size from roughly 2 km^2^ to over 8,000 km^2^ (n=991). It was therefore felt to be prudent to reduce this high spatial variation prior to analyses. To correct for this extreme variation in the spatial resolution of aggregated presence data, in this study, the number of positive LI locations in a given parroquia were reassigned from the centroid of the administrative boundary to cities and villages, using a combination of OpenStreetMap and Google Earth satellite imagery in ArcMap (ver. 10.4) to identify developed areas. This method of informed disaggregation allowed for better spatial representation without compromising de-identification.

### Socio-environmental data acquisition

Environmental coverage datasets for current climatic conditions, comprised of rasterized altitude and 19 derived biophysical variables (Bioclim), were compiled using publicly available interpolated weather station data (WorldClim ver. 1.4., http://worldclim.org) (Table 1) [39]. WorldClim provides long-term climate averages based on weather station records from 1950– 2000, a period coinciding with the start of the MSP’s larval survey. Because *Ae. aegypti* is primarily considered an urban vector in close association with human development, gridded human population density, adjusted to data from the United Nations World Population Prospects 2015 Revision, was also included as an environmental predictor for initial model building as a proxy for built land covers (Socioeconomic Data and Applications Center (SEDAC) Gridded Population of the World (GPW)) [40,41]. A resolution of 2.5 arc-minutes (i.e. 5km grid cells) was chosen for all raster layers to reflect variability in the resolution of geolocated data.

**Table 1.**
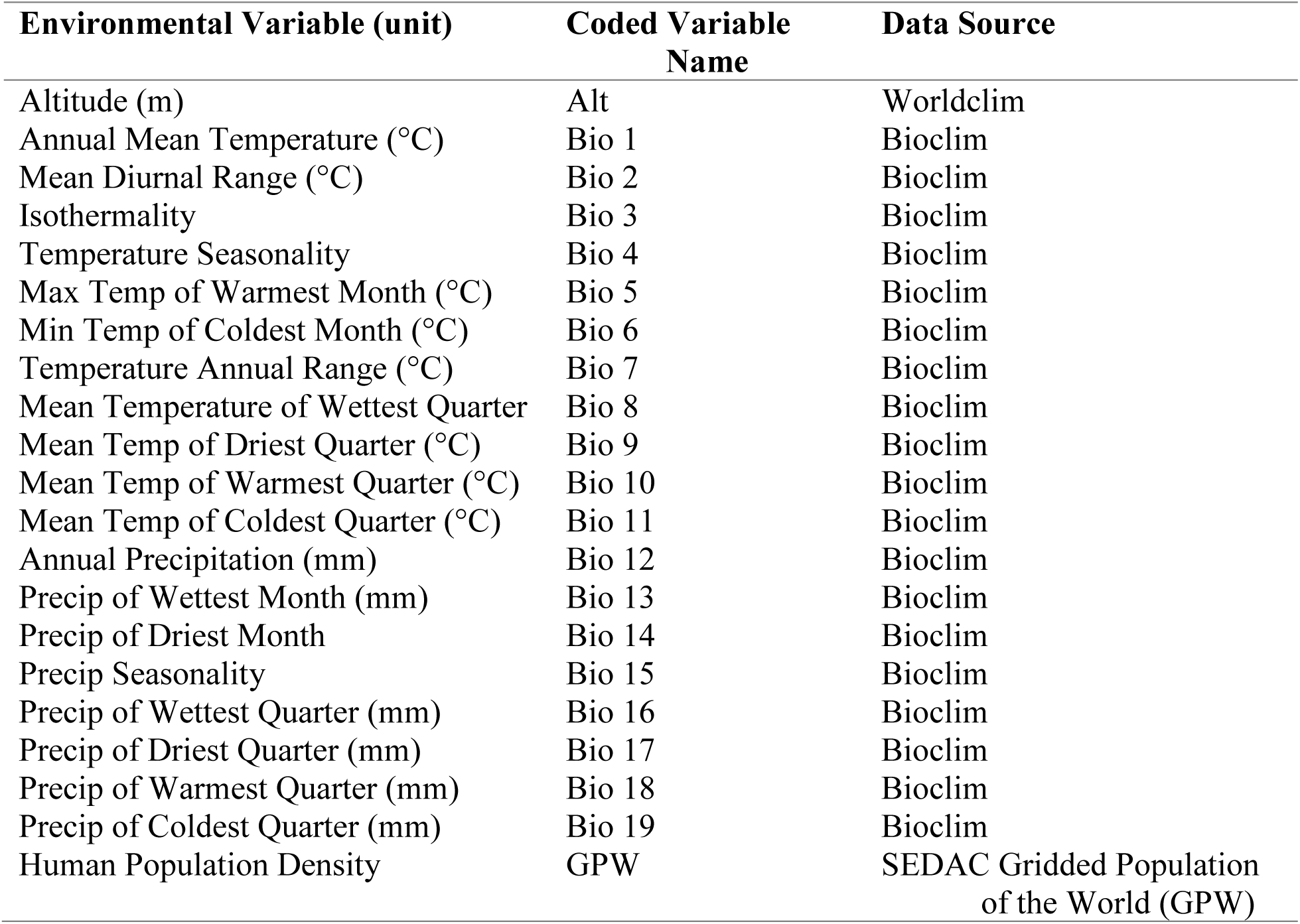
Environmental variables used in building GARP models for *Aedes aegypti* in Ecuador.

| Environmental Variable (unit) | Coded Variable Name | Data Source |
| --- | --- | --- |
| Altitude (m) | Alt | Worldclim |
| Annual Mean Temperature (°C) | Bio 1 | Bioclim |
| Mean Diurnal Range (°C) | Bio 2 | Bioclim |
| Isothermality | Bio 3 | Bioclim |
| Temperature Seasonality | Bio 4 | Bioclim |
| Max Temp of Warmest Month (°C) | Bio 5 | Bioclim |
| Min Temp of Coldest Month (°C) | Bio 6 | Bioclim |
| Temperature Annual Range (°C) | Bio 7 | Bioclim |
| Mean Temperature of Wettest Quarter | Bio 8 | Bioclim |
| Mean Temp of Driest Quarter (°C) | Bio 9 | Bioclim |
| Mean Temp of Warmest Quarter (°C) | Bio 10 | Bioclim |
| Mean Temp of Coldest Quarter (°C) | Bio 11 | Bioclim |
| Annual Precipitation (mm) | Bio 12 | Bioclim |
| Precip of Wettest Month (mm) | Bio 13 | Bioclim |
| Precip of Driest Month | Bio 14 | Bioclim |
| Precip Seasonality | Bio 15 | Bioclim |
| Precip of Wettest Quarter (mm) | Bio 16 | Bioclim |
| Precip of Driest Quarter (mm) | Bio 17 | Bioclim |
| Precip of Warmest Quarter (mm) | Bio 18 | Bioclim |
| Precip of Coldest Quarter (mm) | Bio 19 | Bioclim |
| Human Population Density | GPW | SEDAC Gridded Population of the World (GPW) |

Environmental coverages for estimated future climatic conditions in the year 2050 were taken from forecasted Bioclim variables, allowing for direct comparison between current and future predicted ranges. We chose three general circulation models (GCMs) of physical climate processes commonly used in projecting shifts in species distributions, the Beijing Climate Center Climate System Model (BCC-CSM-1), National Center for Atmospheric Research Community Climate System Model (CCSM4), and the Hadley Centre Global Environment Model version 2, Earth-System configuration (HADGEM2-ES) under the four standard emissions scenarios (RCP 2.6, RCP 4.5, RCP 6.0, RCP 8.5) [23,42–46]. Gridded human population data available through SEDAC are only projected through the year 2020 [40]. To obtain human population for the year 2050, a simple linear extrapolation wherein we assume a stable rate of growth, was performed on a pixel-by-pixel basis in ArcMap with available years of SEDAC data, a growth trend which mirrors more sophisticated cohort-based population estimates for Ecuador projected for the same time period [47,48].

### Ecological niche modeling

Ecological niche models (ENMs) reflecting current and future climate conditions were built using DesktopGARP ver. 1.1.3 (DG) [35]. LI point records and environmental coverage datasets were prepared for modeling using the ‘GARPTools’ package (co-developed by C.G. Haase and J.K. Blackburn) in the program R (ver. 3.3.1). Spatially unique LI records (n=478) were split into 75% training (n=358) and 25% testing datasets (n=119) for ten randomly selected iterations; training datasets were used in model building and testing datasets were used to compute model accuracy metrics [34,35,49]. Ten experiments were run in DG, each using one of the randomly selected LI training datasets and the full set of current environmental coverage variables (Table 2). Each experiment was run for 200 models, allowing for a maximum of 1,000 iterations with a convergence limit of 0.01. Occurrence data were internally partitioned into 75% training/25% testing for model building, and top models were selected using the ‘Best Subsets’ option, specifying a 10% hard omission threshold and 50% commission threshold [50]. The ten top best subsets models from each GARP experiment were summated with the GARPTools package to assess model agreement and accuracy. Model accuracy metrics for each GARP experiment were calculated from the 25% testing dataset withheld from the model building process. Three measures of accuracy, calculated in GARPTools, were used to compare best subsets from each experiment: receiver operator characteristic (ROC) curve with area under the curve (AUC), commission, and omission [51].

**Table 2.** Accuracy metrics for best model subsets built using the full set of environmental coverage variables. Each experiment was performed with a randomly chosen subset (75%) of LI presence points.

| Experiment | AUC | Avg.<br>Commission | Avg. Omission |
| --- | --- | --- | --- |
| 1 | 0.72 | 63.98 | 3.70 |
| 2 | 0.73 | 64.19 | 3.19 |
| 3 | 0.68 | 59.49 | 8.40 |
| 4 | 0.73 | 62.01 | 5.96 |
| 5 | 0.67 | 67.02 | 5.55 |
| 6 | 0.73 | 60.86 | 4.03 |
| 7 | 0.70 | 67.18 | 2.69 |
| 8 | 0.76 | 64.88 | 5.63 |
| **9** | **0.74** | **58.78** | **4.45** |
| 10 | 0.72 | 60.92 | 5.63 |

The model building process was then repeated in DG with the best performing training dataset (i.e. high AUC relative to low omission), comparing full model performance with more parsimonious sets of environmental variables. In addition to variable combinations selected based on previous literature, the GARPTools package was used to extract ruleset trends from the full model (e.g. prevalence and importance of given variables in the resulting model) to assemble additional candidate variable sets for model comparison. The subset of models with the highest AUC and lowest omission (i.e. best model) was chosen as the most probable estimate of current larval mosquito geographic distribution, and rulesets generated from the best model were then projected to the year 2050 for all combinations of GCMs and RCPs. To compare the relative changes in geographic predictions between current climate and future scenarios, the best subsets of current and projected future models for each RCP scenario were recoded as binary geographic distributions (i.e. presence and absence) in ArcMap, where cells with model agreement of ≥ 6 were considered present. Recoded distributions were combined using the ‘Raster Calculator’ tool in the Spatial Analyst extension of the program ArcMap, allowing for the visualization of range agreement across GCMs. The number of people at risk in areas of expanding mosquito distribution, where range expansion was predicted under at least one GCM, was estimated in ArcMap, using the Raster Calculator tool to extract information on GPW and extrapolated population for the year 2050.

## Results

The original dataset of LI occurrences in Ecuador, provided by the MSP, consisted of 3,655 collection events aggregated to 374 parroquia centroids, indicating the number of parroquias that had positive surveillance results for *Ae. aegypti* larvae during the study period. Dis-aggregation of these data yielded 478 spatially unique locations within these parroquias, corresponding with areas of human habitation regularly surveyed by the MSP. Incorporating prior knowledge regarding the agency’s collection of data in developed areas allowed for the adoption of a finer spatial scale for analysis without changing the overall distribution of larval mosquito presence in Ecuador (e.g. mosquitoes remained conspicuously absent in most high-elevation parroquias located in the Andes mountains).

Much of Ecuador was predicted to be suitable for the presence of *Aedes aegypti* larvae under current climatic conditions, with the notable exception of the eastern portion of the country associated with the Amazon basin and high elevation areas associated with the Andes mountain range, running north to south through the center of the country (Fig 2). This iteration of model subsets generated by GARP had the highest AUC, relative to low omission (AUC=0.73, Avg. Commission=63.47, Avg. Omission=3.02), and was built with a reduced set of environmental variables including altitude, human population, maximum temperature of the warmest month, temperature annual range, mean temperature of the wettest month, mean temperature of the driest month, mean temperature of the warmest quarter, mean temperature of the coldest quarter, precipitation of the wettest month, precipitation seasonality, precipitation of the driest quarter, and precipitation of the coldest quarter (Table 3).

**Table 3.**
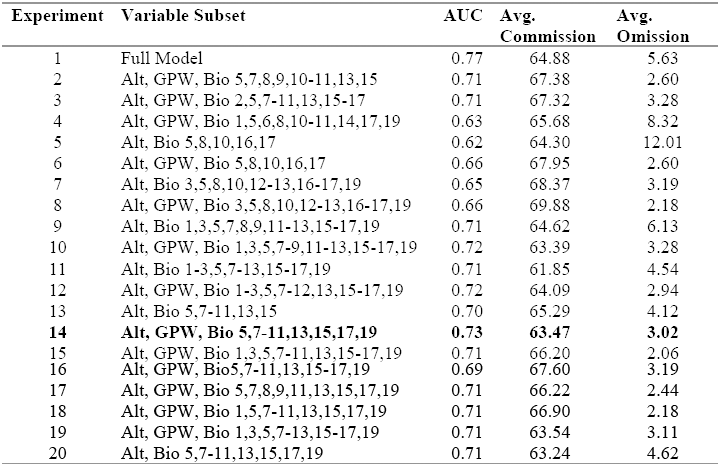
Accuracy metrics for best model subsets built using the best-ranked training dataset and selected subsets of environmental coverages.

**Fig. 2.**
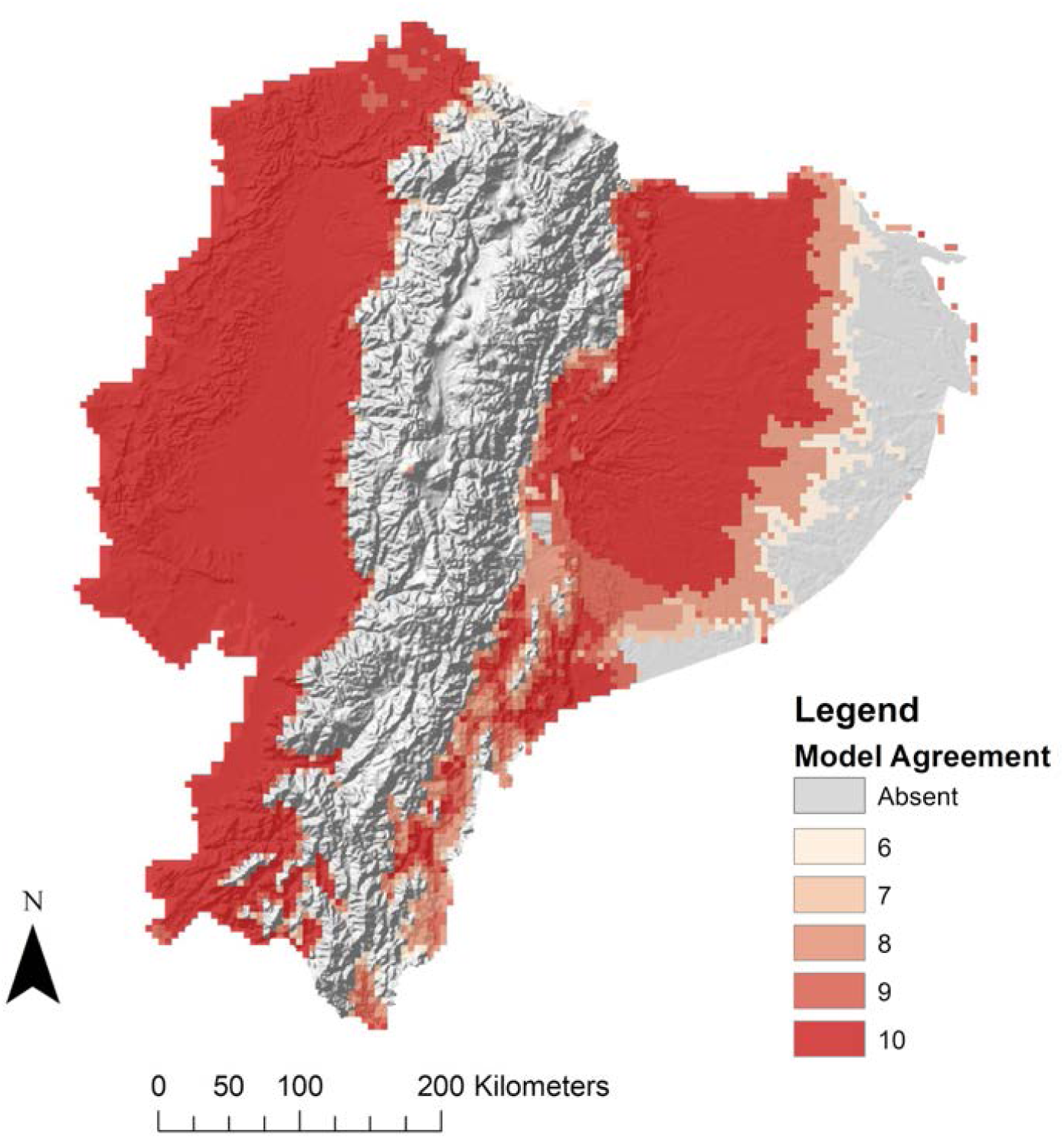
Agreement of best model subsets built with best-ranked suite of environmental variables for larval *Aedes aegypti* presence in Ecuador under current climate conditions. Models had high levels of agreement in the western coastal lowlands, and lower levels of agreement in the eastern Amazon basin.

The projected geographic distribution of larval *Ae. aegypti* for the year 2050 (Fig 3B1 and 3B2, 3C1 and 3C2, 3D1 and 3D2, S1 and S2 Figs), built with the best-performing selection of environmental coverages under four climate change scenarios, showed marked changes in pattern when compared with estimated mosquito presence under current conditions (Fig 3A1 and 3A2, S1 and S2 Figs). Potential distributional shifts were generally consistent across GCMs, with slight range expansions into areas of higher elevation and large portions of the eastern Amazonian basin predicting mosquito absence (Fig 3, S1 and S2 Figs). Combining the current and future model agreement rasters for best subset models by RCP revealed predicted areas of geographic stability in western Ecuador and the eastern foothills of the Andes, range contraction throughout much of Amazon basin in the east, and range expansions along transitional elevation boundaries over time (Fig 4). Range expansions and contractions were generally consistent across climate models, with the magnitude of distribution change increasing with more extreme climate change scenarios (Fig 4). Similarly, the human population with the potential to experience increased exposure to mosquito presence generally increases with RCP, with an additional 9,473 (RCP2.6), 11,155 (RCP4.5), 10,492 (RCP6.0), and 12,939 (RCP8.5) people currently living in areas of transitional elevation estimated at risk of becoming exposed under different climate change scenarios (Table 4).

**Table 4.** Estimated human population inhabiting areas of transitional elevation in Ecuador, which may experience increased exposure to moquito-borne disease transmission under climate change.

| **Representative**<br>**Concentration**<br>**Pathway (RCP)** | **GPW 2010**<br>**Population** | **Projected 2050**<br>**Population** | **Area (km<sup>2</sup>)** |
| --- | --- | --- | --- |
| **RCP 2.6** | 9,473 | 15,399 | 2,755 |
| **RCP 4.5** | 11,155 | 18,439 | 3,530 |
| **RCP 6.0** | 10,492 | 17,100 | 3,155 |
| **RCP 8.5** | 12,939 | 21,298 | 4,215 |

**Fig. 3.**
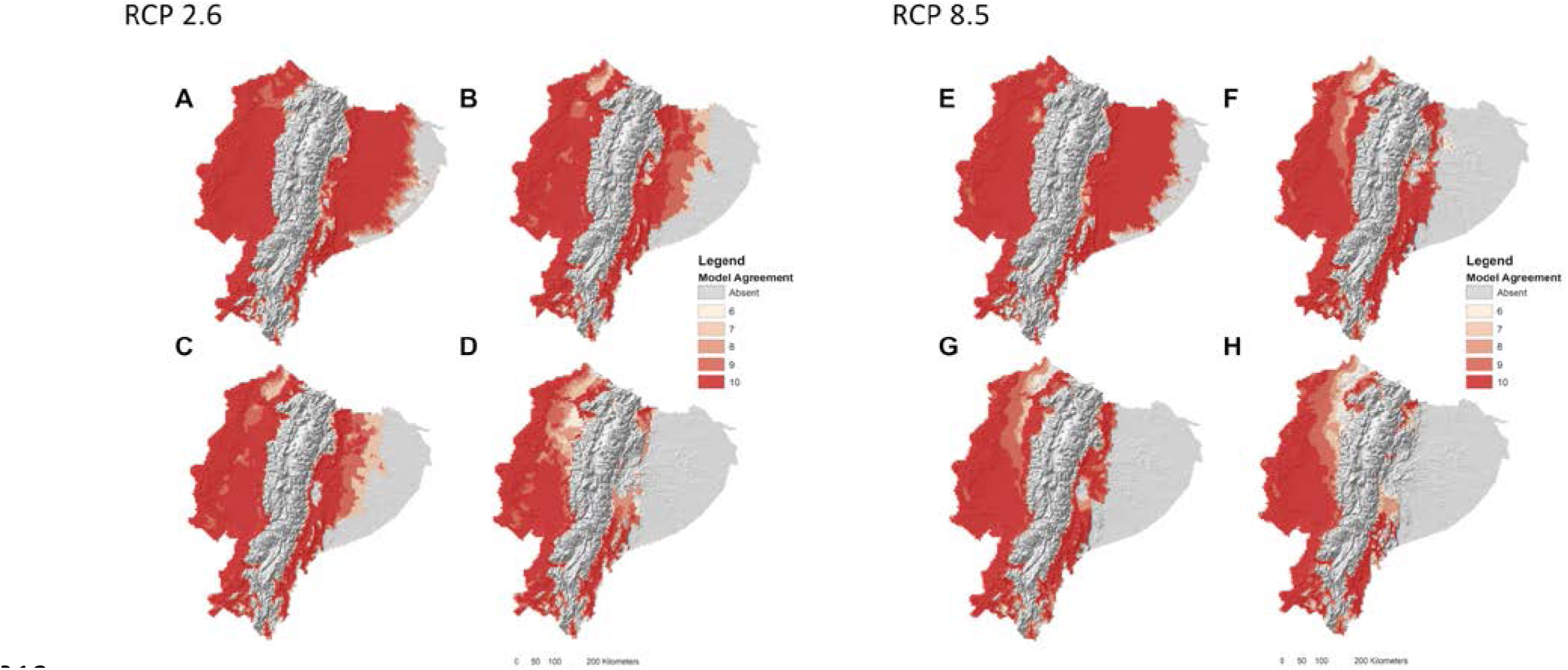
Agreement of best model subsets built with best ranked suite of environmental variables for larval *Aedes aegypti* presence in Ecuador under A) current climate conditions and future climate conditions projected to the year 2050 under Representative Concentration Pathway (RCP) 2.6 (B1,C1,D1) and 8.5 (B2,C2,D2) for the B) BCC-CSM-1, C) CCSM4, and D) HADGEM2-ES General Circulation Models (GCM) climate models.

**Fig. 4.**
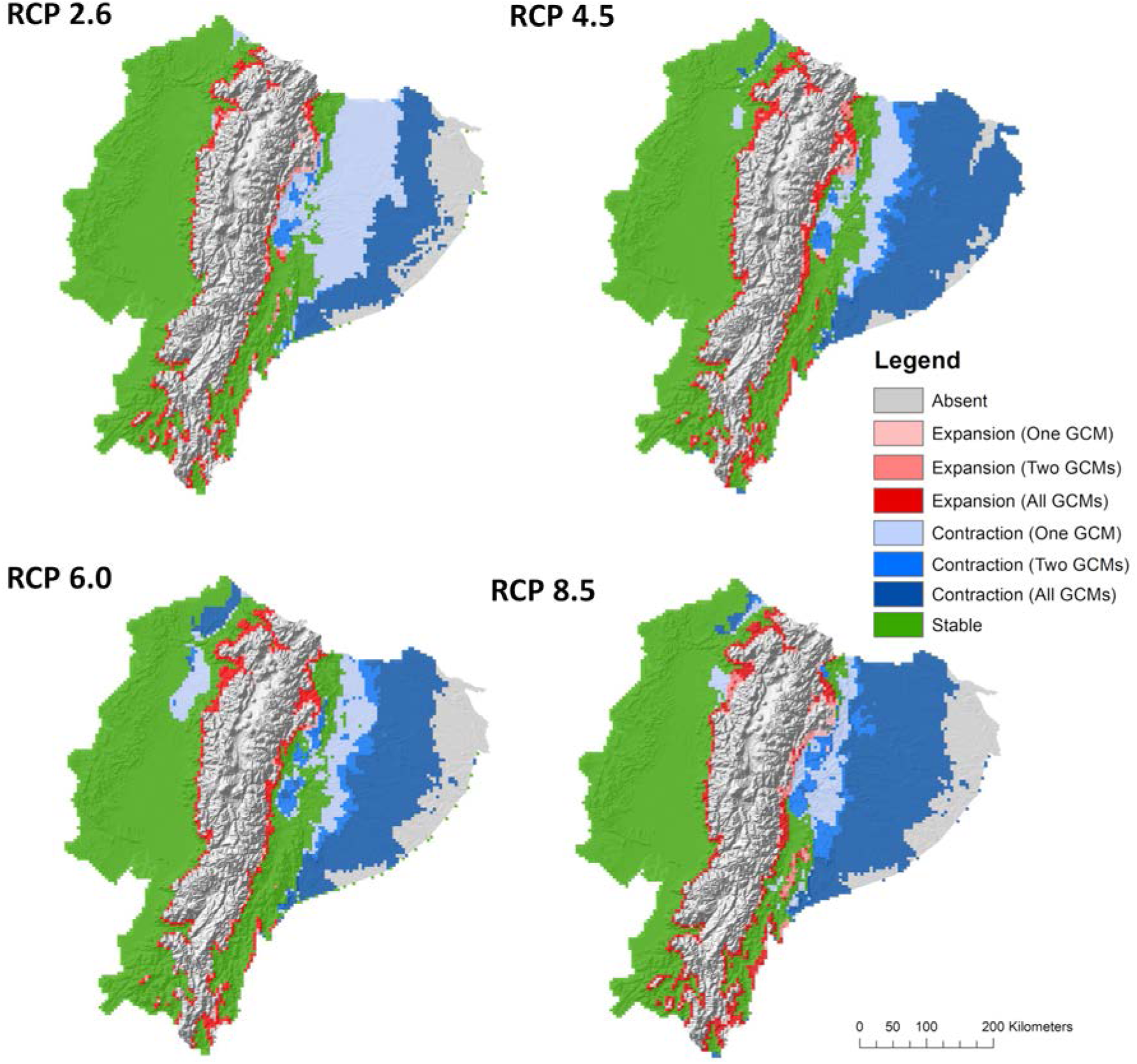
Best model subsets for current and future climate (GCMs projected to the year 2050) were combined by RCP emissions scenarios to illustrate the estimated contraction and expansion of larval *Aedes aegypti* geographic range in Ecuador.

## Discussion

The predicted current geographic distribution of larval *Aedes aegypti* suitability in Ecuador, under current climate conditions, largely reflects present-day risk maps for many of the mosquito-borne diseases currently circulating in the country, wherein populations living at high altitudes are not considered at-risk for transmission [52]. Predicted larval distributions are roughly continuous in the eastern and western portions of Ecuador, but are sharply restricted along increasing elevation gradients in the central portion of the country, the area corresponding with the location of the Andes mountain range (Fig 2) [9]. This conspicuous absence of mosquitoes in the Andes reflects the generally protective nature of high mountain elevations from mosquito presence, with all models predicting larval mosquito absence throughout central Ecuador (Figs 2–6). The predicted absence of *Ae. aegypti* in the eastern portion of the Amazon basin is notable, as this is a region currently perceived as high-risk for mosquito exposure by public health officials despite having low human population density, mostly owing to its low altitude (Fig 2). Although similar in elevation to regions of active disease transmission in the West, the hydrology of the Amazon basin differs considerably from coastal areas. Previous work in this region suggests a great deal of spatial variability in the basin with regards to precipitation and drainage patterns [53,54]. Given that the mosquito life cycle depends heavily on the availability of water in the environment, spatial discrepancies in precipitation could account for the low model agreement of mosquito presence in the easternmost portion of the Amazon.

Range expansion of *Ae. aegypti* into higher elevations as a result of changing climate was supported across GCM models and emissions scenarios (Figs 3–7). All best model subsets suggest that areas of transitional elevation along the eastern and western peripheries of the Andes mountains may experience some level of increased exposure to the presence of mosquitoes, though much of the mountain range, including densely populated areas like the capital city, Quito, will remain unsuitable habitat. The intrusion of *Ae. aegypti* into areas of transitioning elevation represents a potential area of concern for public health managers, as communities in these areas are largely protected from mosquito exposure and associated diseases under current climatic conditions. Excluding travel-related cases, reporting of arboviral diseases in Ecuador’s mountain dwelling populations is quite low, although there are low-lying valleys near Quito that may be suitable for arbovirus transmission. Accordingly, the MSP primarily directs mosquito-borne disease outreach and intervention efforts to high-risk communities, particularly in large coastal cities with consistently high disease incidence, such as Guayaquil and Machala. As a result, communities situated in the foothills of the Andes will not necessarily have the same risk perceptions and preventative behaviors as those communities burdened with historically high incidence of mosquito-borne diseases. This sets the stage for potential disparities in preventative knowledge and health services should *Aedes* mosquitoes expand into naïve populations [5,55]. Conversely, extirpation of *Ae. aegypti*, especially the large range contraction predicted in Amazonian Ecuador, has the potential to conserve valuable resources by triggering allocation shifts as unsuitable areas no longer support active disease transmission.

Our findings are broadly consistent with a previous coarser scale ENM analysis of adult mosquitoes in Ecuador, which suggests that while *Aedes* mosquitoes may shift into highland areas under changing climate conditions, the total area of suitable habitat will ultimately decrease as localized climatic conditions favor extirpation [30]. However, models of *Aedes* distribution in the previous study were made through the year 2100, representing an extended time horizon for guiding agency decision making. While predicted ranges in 2100 are visually similar to results presented here, notable discrepancies exist between the spatial distributions predicted in our models and the previous study for 2050, where the previous model predicts widespread absence of mosquitoes in central Ecuador and presence throughout much of the eastern Amazon basin. In contrast to our methods, Escobar *et al.* [30] used a different niche modeling algorithm, a different model of climate change (A2), a coarser spatial resolution (20 km), and combined global species occurrence for two adult arbovirus vectors, *Ae. aegypti* and the Asian tiger mosquito (*Aedes albopictus*), to predict pooled arbovirus risk throughout Ecuador. Though *Ae. aegpyti* and *Ae. albopictus* are competent vectors of diseases that occur in Ecuador (e.g. dengue, chikungunya, Zika), these species differ significantly in their physiology, possibly driving observed discrepancies between the models of pooled adult *Aedes spp.* risk and larval *Ae. aegypti* range [56]. Reaching consensus across ENMs is a known area of conflict in ecology that requires more research, where various methodologies can lead to vastly different forecasts of geographic distributions and risk, making direct comparisons between models difficult [57]. Future studies combining multiple approaches and comparing the impact of input on models could help resolve this conundrum.

We chose a moderately low spatial resolution for this study (5km raster cells) to reflect the highest level of precision that could be assigned to larval mosquito occurrence (i.e. points could be matched to cities or clusters of villages, but not to individual households or neighborhoods). This scale of analysis presents a limitation in applying resulting ENMs for local management decisions. Arboviral disease transmission and larval mosquito presence, especially for *Ae. aegypti*, are typically managed at the household or neighborhood level, and although we can use these results to discuss regional changes in mosquito distribution throughout Ecuador, we cannot overstate the findings as a means to assess risk at the level of disease transmission [58]. Furthermore, the LI survey conducted by the MSP was limited in that focus was placed on sampling areas with perceived arbovirus transmission risk throughout Ecuador, especially households in densely populated urban centers and established communities where cases had been reported in the past. Low accessibility and human population density in Ecuador’s eastern basin region may have contributed to under sampling of mosquito presence in these areas, possibly accounting for low model agreement in this area. Ultimately, robust vector surveillance for *Ae. aegypti* in eastern Ecuador would be required to validate absence in this region, though such intensive ground-truthing would be wrought with logistical concerns, including diversion of scarce surveillance resources from high-demand management districts and the inherent difficulty of establishing “true” absence via surveys.

*Aedes aegypti* is a globally invasive species, owing much of its success to its close connection with human activity and urban environments. As a result, microclimate can become a critical factor in determining true habitat suitability, and there are many examples of anthropogenic structures providing a buffering effect, or refuge, against climatic conditions that would be otherwise physiologically limiting to insect vectors [5,59–62]. Similarly, dramatic shifts in species compositions in Ecuador, mediated by elevation, also occur on very fine spatial scales [63,64]. Moving forward, observed areas of range expansion on the edge of unsuitable habitat may be better modeled at finer resolutions, which would aid in making community-targeted management decisions based on estimated risk.

Based on the results of this study, we conclude that the geographic distribution of larval *Aedes aegypti* in Ecuador will be impacted by projected shifts in climate. Extensive changes in modeled vector distributions were observed even under the most conservative climate change scenario, and these changes, although consistent in pattern, became more evident with increasingly high greenhouse gas emissions scenarios. Although there is a continued need for surveillance activities, these findings enable us to anticipate transitioning risk of arboviral diseases in a spatial context throughout Ecuador, allowing for long-term planning of agency vector control strategies.

## Acknowledgements

Many thanks to the field technicians and other staff at the National Service for the Control of Vector Borne Diseases of the Ministry of Health of Ecuador, who generated the data used in this analysis.

## Supporting information

**S1 Fig.**
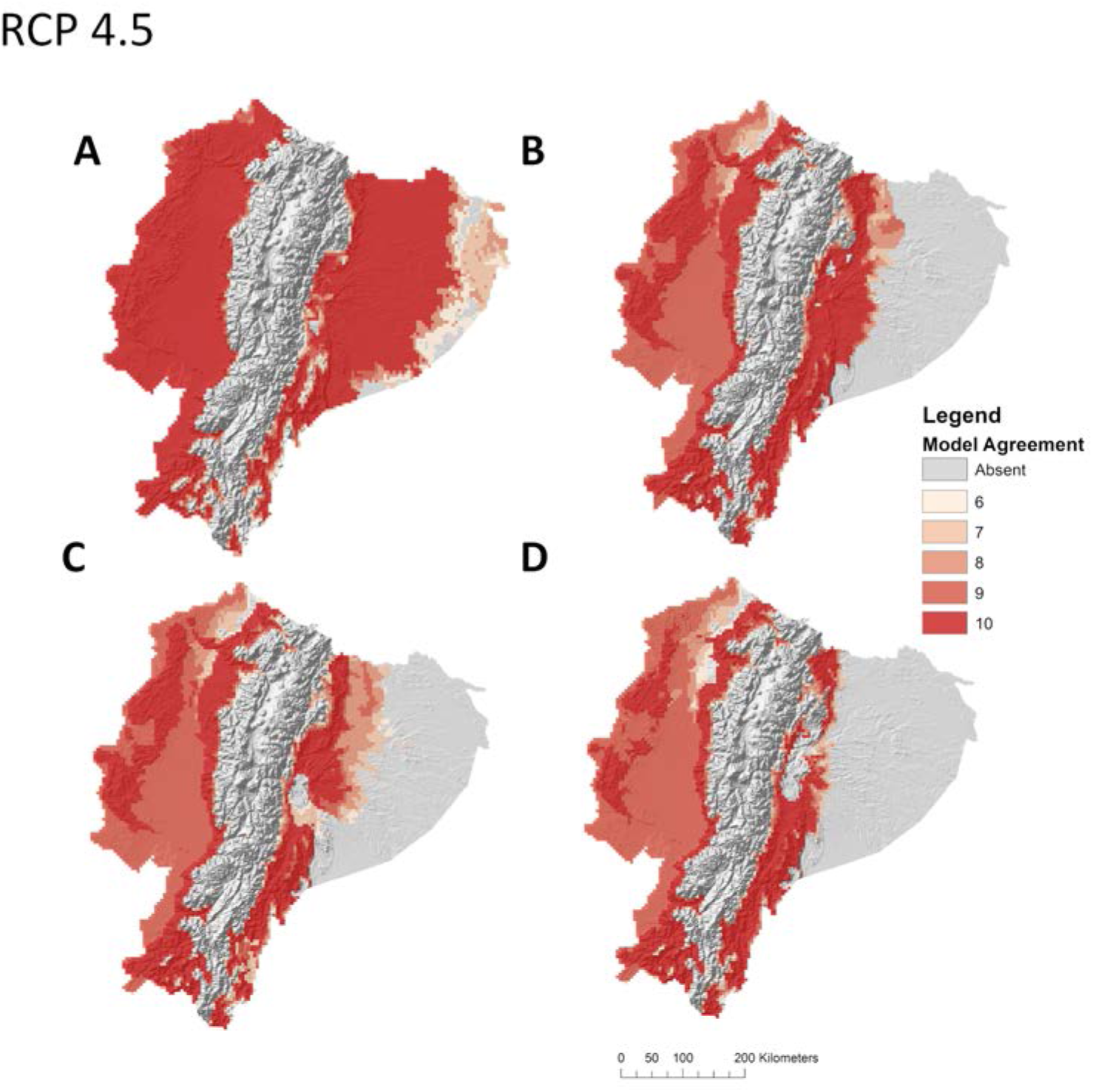
Agreement of best model subsets built with best ranked suite of environmental variables for larval *Aedes aegypti* presence in Ecuador under A) current climate conditions and future climate conditions projected to the year 2050 under Representative Concentration Pathway (RCP) 4.5 for the B) BCC-CSM-1, C) CCSM4, and D) HADGEM2-ES General Circulation Models (GCM) climate models.

**S2 Fig.**
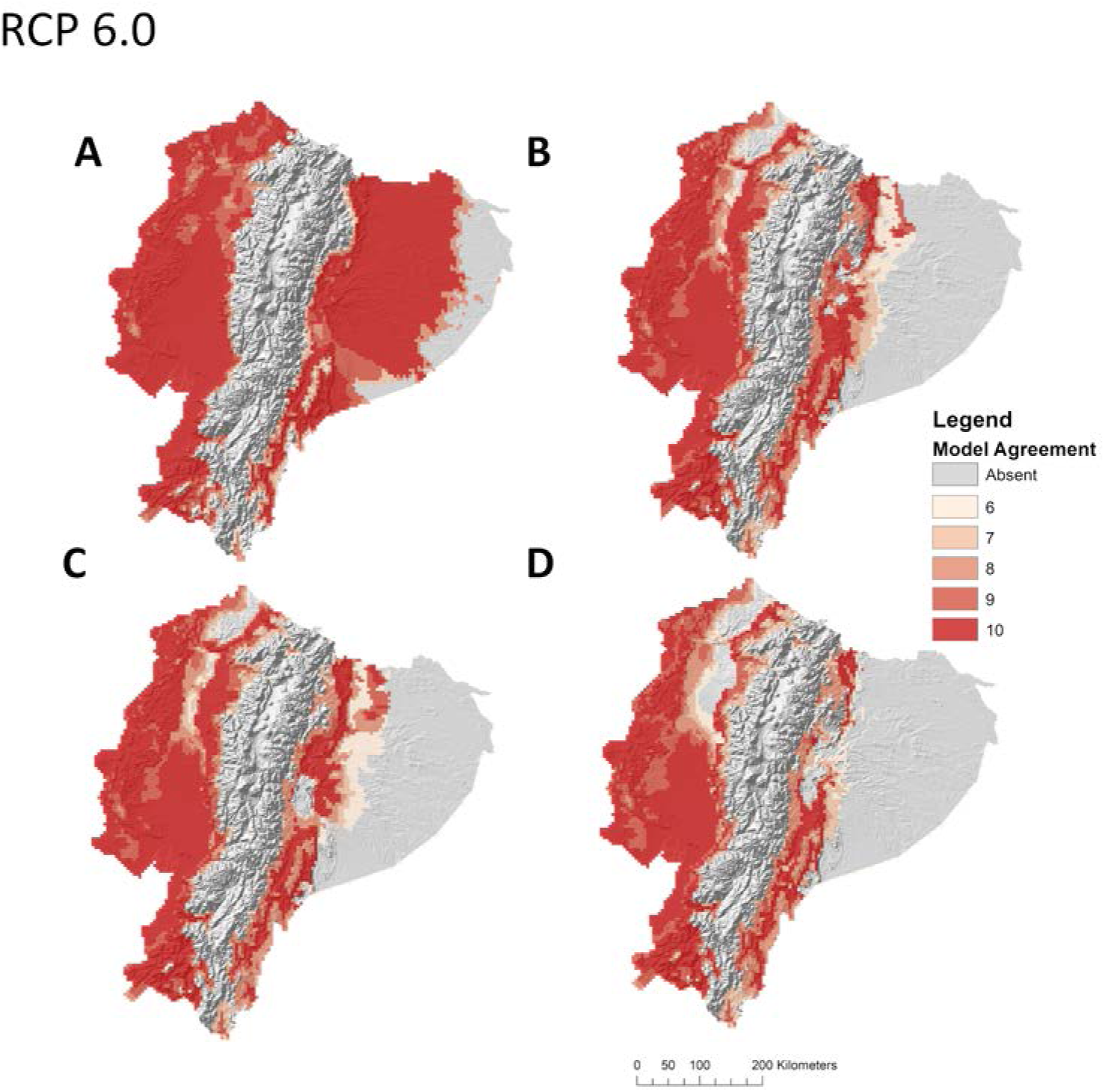
Agreement of best model subsets built with best ranked suite of environmental variables for larval *Aedes aegypti* presence in Ecuador under A) current climate conditions and future climate conditions projected to the year 2050 under Representative Concentration Pathway (RCP) 6.0 for the B) BCC-CSM-1, C) CCSM4, and D) HADGEM2-ES General Circulation Models (GCM) climate models.

**S1 Table.** Prevalence of environmental coverages in model building ruleset.

| Environmental Variable | Ruleset Prevalence |
| --- | --- |
| Alt | 0.94 |
| Bio 5 | 0.94 |
| Bio 7 | 0.91 |
| Bio 8 | 0.82 |
| Bio 9 | 0.85 |
| Bio 10 | 0.85 |
| Bio 11 | 0.74 |
| Bio 13 | 0.88 |
| Bio 15 | 0.68 |
| Bio 17 | 0.94 |
| Bio 19 | 0.85 |
| GPW | 0.74 |

